# Different mycorrhizal nutrient acquisition strategies shape tree species competition and coexistence dynamics

**DOI:** 10.1101/2021.10.26.465925

**Authors:** Michael E. Van Nuland, Po-Ju Ke, Joe Wan, Kabir G. Peay

**Affiliations:** Department of Biology, Stanford University, Stanford, USA; Institute of Ecology and Evolutionary Biology, National Taiwan University, Taipei, Taiwan; Institute of Integrative Biology, Department of Environmental Systems Science, ETH Zürich, Zürich, Switzerland

**Author notes:** **Statement of authorship**: MVN and KGP designed the experiment. MVN created the experiment, collected data, conducted the analyses, and wrote the initial manuscript draft. PJK and JW helped interpret the results. All authors contributed significant manuscript edits. **Data accessibility statement**: All data and code will be archived on Zenodo prior to publication.

**Keywords:** arbuscular mycorrhizal fungi, coexistence, competition, ectomycorrhizal fungi, nitrogen, plant-soil feedback, soil nutrients

## Abstract

Mycorrhizal fungi with different nutrient acquisition strategies influence the functional separation among plant species. This might drive resource competition dynamics that cumulatively impact tree species coexistence, but few manipulative experiments have directly tested this. Combining surveys and experiments in a modern coexistence theory framework, we tested how variation in mycorrhizal strategies and nutrient conditions affect competitive outcomes between co-occurring ectomycorrhizal (EM) and arbuscular mycorrhizal (AM) tree species. The dependency on EM symbioses increased with latitude and nitrogen (N) limitation, matching global trends. Host-specific soil microbiome conditioning and N fertilization combined to qualitatively affect coexistence outcomes. Lower N conditions favored EM over AM trees, but N fertilization reversed this outcome for southern species, consistent with regional-scale forest mycorrhizal transitions. As the magnitude and outcome of microbially-mediated competition depends on mycorrhizal differentiation and soil nutrient availability, this strongly supports the importance of mycorrhizal symbioses in driving large-scale biogeographic patterns of tree species.

## Introduction

The soil microbiome plays an important role in shaping plant species coexistence and community diversity as plants build microbial relationships that influence host plant performance, competitive ability, and functional separation among co-occurring species (Peay 2016; Kandlikar 2019; Ke & Wan 2020). Mycorrhizal fungi are of particular importance as these root-associated microbial symbionts influence ecosystem dynamics affecting nutrient availability and plant productivity (van der Heijden et al. 2015). Nearly all tree species on Earth form symbioses with one of two main mycorrhizal types (Brundrett 2009; Steidinger et al. 2019) that generally appear to have different nutrient acquisition capabilities: arbuscular mycorrhizal (AM) and ectomycorrhizal (EM) fungi. Although nutrient uptake strategies can vary within these diverse assemblages (Lilleskov et al. 2002; Maherali & Klironomos 2007), AM fungi are generally better at scavenging soil phosphorus (P) and inorganic nutrients, while EM fungi use specialized extracellular enzymes for acquiring nutrients – especially nitrogen (N) – from complex organic sources (Bodeker et al. 2009; Tisserant et al. 2013; Lindahl & Tunlid 2015; Kohler et al. 2015). These mycorrhizal-mediated nutrient uptake patterns might influence the competitive balance between plant species with different mycorrhizal types (Aerts 2003; Peay 2016), but there has been little experimental confirmation of such potentially widespread ecological processes affecting species coexistence.

Plant communities with competing species are stabilized when within-species competition is stronger than among-species competition (Adler et al. 2018). Competitive outcomes are determined by the amount of niche differences between competitors (e.g., the differences in species’ requirements and use of limiting resources) as well as underlying fitness inequalities (e.g., competitive hierarchies arising from growth rate or reproductive differences). Modern coexistence theory describes how coexistence can be achieved based on stabilizing and equalizing mechanisms (Chesson 2000). Specifically, conditions favoring species coexistence occur when stabilizing mechanisms increase species niche differences (or reduce niche overlap), or when equalizing mechanisms act to reduce fitness differences between competitors. Modern coexistence theory is an extremely promising framework because it provides a way to quantitatively link different facets of plant ecology with their ultimate goal of understanding how and why plant species coexist (Adler et al. 2006; Grainger et al. 2019; Johnson 2021).

While modern coexistence theory continues to develop, its applications to plant ecology have been largely isolated from ecological interactions with soil microbes. By contrast, the reciprocal interactions between plants and soil microbes, and their role in biodiversity maintenance, have been more extensively studied under the framework of plant-soil feedback (van der Putten et al. 2013). Plant-soil feedbacks arise when plants experience net costs or benefits from interacting with their host-specific soil biota compared to a competitor’s soil community (Bever 2003). Negative feedbacks stabilize community diversity and promote species coexistence through negative frequency-dependent plant population dynamics (favoring species when rare), while positive feedbacks destabilize community dynamics and can lead to monodominance (Bever et al. 1997; Eppinga et al. 2018). Plant responsiveness to various soil microbial groups (e.g., pathogens and mutualists) can drive feedbacks in different directions, and a recent study of North American trees found that negative and positive feedbacks are associated with trees that form AM vs. EM fungal symbioses, respectively (Bennett et al. 2017). However, plant-soil feedback experiments rarely account for plant-plant competition directly which makes it difficult to predict the long-term competitive outcomes from specific coexistence mechanisms (Lekberg et al. 2018; Ke & Wan 2020). Recent theoretical studies have proposed experimental designs that combine elements from both modern coexistence theory and plant-soil feedback studies, making it possible to address this knowledge gap by incorporating soil microbial effects and plant-plant competition into the calculations of niche and fitness differences (Kandlikar et al. 2019; Ke & Wan 2020). This opens the door to resolve long-standing hypotheses about how host-specific differences in mycorrhizal associations and soil microbiome conditioning modify plant-plant interactions and species distribution patterns.

One long-standing hypothesis is that competitive outcomes between AM and EM tree species are determined by the identity and form of major limiting nutrients (Aerts 2003; Peay 2016). At the global scale, AM tree species are more prevalent near the equator while EM tree species become progressively more dominant at higher latitudes, a transition which corresponds with a climatically driven shift from fast cycling, N rich soils at low latitudes to slow cycling, N-limited soils at high latitudes (Read 1991; Steidinger et al. 2019; Du et al. 2020). While it is generally accepted that regional-scale dominance of specific mycorrhizal symbioses is driven by adaptation to different soil nutrient environments–inorganic N for AM and organic N for EM trees and fungi (Lu & Hedin 2019)–the assumed competitive mechanisms have not been directly tested. These competitive mechanisms center on the mycorrhizal-mediated changes in niche and fitness differences that may drive competitive exclusions depending on whether AM or EM associations are favored under resource competition. In addition, most studies of mycorrhizal biogeography assume that all trees within an AM or EM functional guild participate equally in their respective fungal symbiosis, even though the degree of mycorrhizal association can vary among and within plant species across geographic regions (Gehring et al. 2006; Soudzilovskaia et al. 2015; Karst et al. 2021a; Bueno et al. 2021).

In this study we use the framework of modern coexistence theory to test how nutrient availability and symbiont dependency affect the outcome of competition between trees with different mycorrhizal strategies. We manipulated competitive environments under different nutrient regimes between EM and AM tree genera (*Populus* vs. *Acer*) and congeneric species with differing geographic and climatic ranges (*P. tremuloides* and *P. deltoides*). We hypothesized that:

1. Dependency on EM symbioses increases with latitude and N limitation, such that EM root colonization is greater for tree species with more northern geographic distributions and decreases with mineral N fertilization.
2. The strength and direction of plant-soil feedbacks differ among plant mycorrhizal types, with AM trees creating negative feedbacks and EM trees creating positive feedbacks.
3. The competitive advantages of AM and EM associations depend on soil N availability such that EM tree species gain an advantage under low N (varying with EM dependency levels) and mineral N fertilization tips the competitive balance in favor of AM over EM symbioses.

Overall, we show that mycorrhizal type, dependency, and nutrient availability interactively drive plant competition and coexistence outcomes that match host species distributions and global transitions in forest symbioses.

## Methods

### Sample collection and growth room competition experiment

In the field survey, we examined mycorrhizal affinity in two widespread *Populus* species (which are dual mycorrhizal but predominately form EM symbioses in mature trees) that have latitudinally divergent species ranges (Figure 1A). In Summer 2018, we collected *P. deltoides* roots from 24 sites covering Mississippi to Minnesota, and *P. tremuloides* roots from 13 sites in the Rocky Mountains and upper Midwest. At each site we collected ∼12 cm long root sections from five separate trees (5-7 samples/tree), which were gently washed before measuring percent root length colonized by EM fungi using the gridline intersect method (Giovannetti & Mosse 1980). We extracted mean annual temperature and annual precipitation from GPS coordinates taken at the center of each site (WorldClim database; Frick & Hijmans 2017) which were used to estimate site-level decomposition coefficients predicted by the Yasso07 model of climate controls on mass-loss rates of different leaf litter nutrient pools (Tuomi et al. 2009). These decomposition constants have been the best predictors of mycorrhizal dominance in tree communities (Steidinger et al. 2019) as they approximate the speed with which nutrients move from organic matter into more easily accessible mineral forms, and thus whether systems are “fast” or “slow” regarding nutrient cycles.

**Figure 1.**
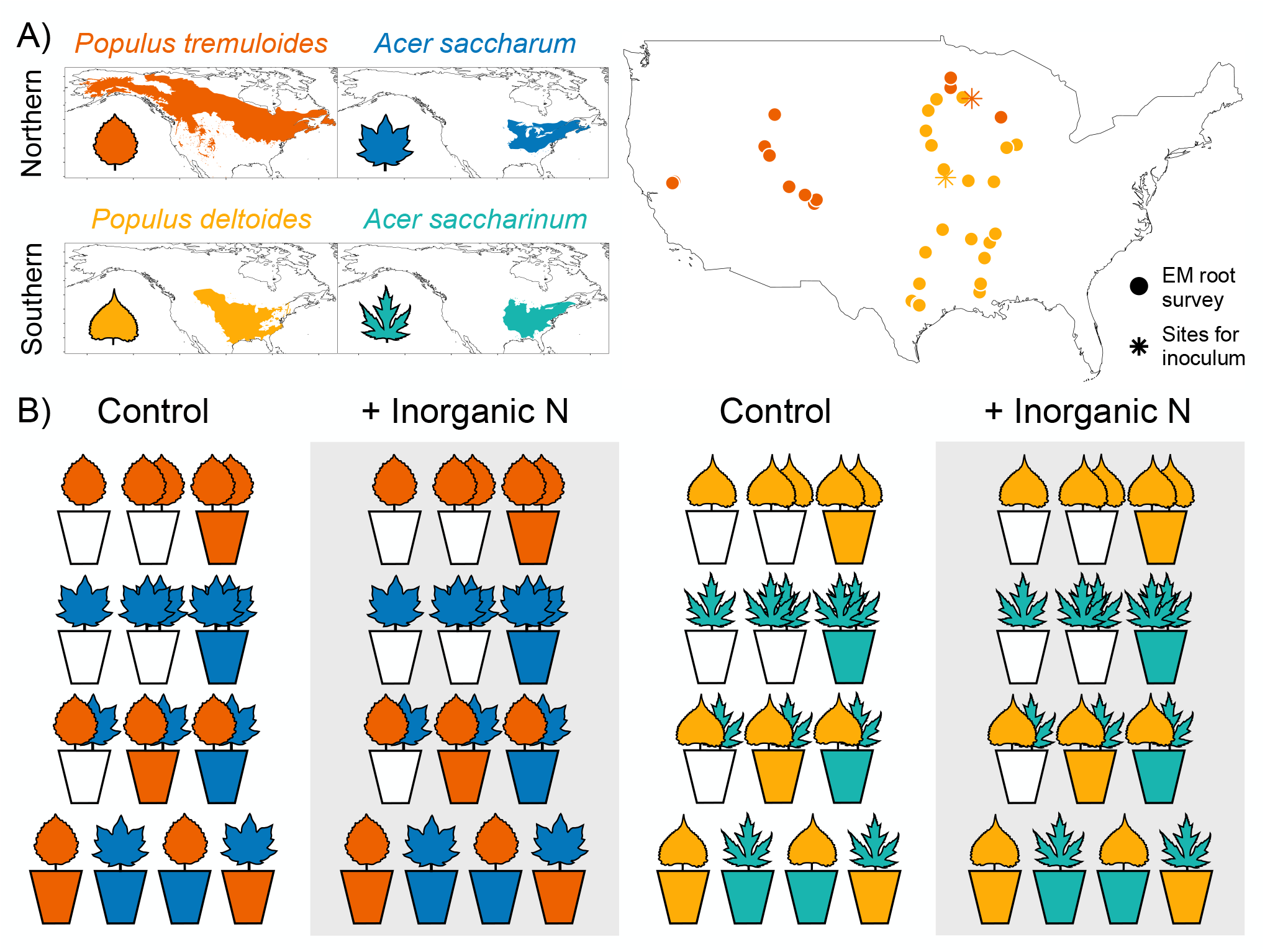
Tree species range maps, sampling locations, and experimental design. **(A)** Range maps of the *Populus* and *Acer* species included in this study, grouped into Northern and Southern pairs. Locations of sites across the US where *Populus* roots were collected for ectomycorrhizal measurements (circles) and soil/root samples were collected for experimental inoculum (stars). **(B)** Outline of the experimental design with soil inoculum, plant competition, and nitrogen fertilization treatments applied to the Northern and Southern tree species pairs. This design was replicated without pooling soil inoculum. Colored pots represent Live inoculum and empty pots show Sterilized inoculum that serve as uncultivated “reference” soil treatments. The first two rows are used to calculate intraspecific effects, the third row is used for interspecific effects, and the final row comprises treatments for plant-soil feedback measurements.

For the growth room experiment, we sampled from the same *Populus* and two *Acer* species that form AM symbioses and generally have northern vs. southern distributions: *A. saccharum* and *A. saccharinum* (Figure 1A). In early Fall 2019, we re-sampled soils and roots beneath trees from two previously visited sites: one northern site where *P. tremuloides* and *A. saccharum* co-occur (elevation=370 m; mean temperature=6.6° C; annual precipitation=802 mm; soil pH=6.9, soil organic matter=4.6% rating), and one southern site where *P. deltoides* and *A. saccharinum* co-occur (elevation=221 m; mean temperature=12.9° C; annual precipitation=956 mm; soil pH=7.4; soil organic matter=3.1% rating). Five pairs of *Populus* and *Acer* trees were sampled at each of the sites (ten total trees per site), with each pair serving as an experimental replicate (i.e., samples were not pooled for the inoculum treatments; Reinhart & Rinella 2016). We collected enough field soil and fine root material to be used for Live and Sterilized (two 45-minute autoclave cycles with a 12hr rest period in between) inoculum treatments, as well as sterilized background soil that we purposefully enriched with sterilized plant leaf material to create a baseline of enriched soil organic nutrient sources (see Supplemental Information for more details).

To test how plant-soil and plant-plant interactions modify tree species competitive outcomes we created a manipulative experiment based on a design that efficiently measures the impact of soil microbes on plant competition and coexistence (Ke & Wan 2020). For each geographic pair of *Populus* (EM) and *Acer* (AM) species, plants were first grown either alone (single individual), in intraspecific competition (two individuals of the same species), or in interspecific competition (one focal individual plus a competitor belonging to the other species) with Sterilized inoculum. For the competition pairs, we also included additional treatments where Live soil cultivated by the competitor were added as inoculum. For example, when the focal *Acer* species was growing in interspecific competition treatments with *Populus*, soils conditioned by the *Populus* species were added. This allowed us to quantify the combined impact of a competitor individual through both plant-plant interaction and plant-soil interaction pathways. We also included four additional treatments with each tree species grown alone with Live inoculum in a reciprocal Home and Away plant-soil feedback design (Bever et al. 2010). This design (in total 13 planting combinations; Figure 1B) was replicated five times by using the separate inoculum sources from each of the five pairs of *Populus* and *Acer* trees sampled at each site. Additionally, we replicated all competition and soil inoculum treatments across two soil nutrient conditions, unfertilized control treatments and mineral N fertilization with NH_4_NO_3_ (5 mL doses at 20 mg N kg^-1^ soil), which we confirmed using Plant Root Simulator resin membrane probes (see Supplemental Information for more details).

We used a randomized block design (one block for each soil inoculum replicate per N fertilization treatment) with plants grown in racks on different shelves in the growth chamber. Plants were transferred into the experiment at approximately the same size, and soil inoculum was applied as ∼5% total soil volume (consistent with other plant-soil feedback designs, Crawford et al. 2019; see Supplemental Information for more details). The experiment lasted 35 weeks, after which harvesting occurred by clipping plants at their base and drying the shoots at 60°C for 72 hours before weighing the aboveground biomass (mg). We gently rinsed roots to remove soil particles and stored root samples in DI water at 4°C for up to 48 hours. EM colonization was measured on *Populus* roots (as above) on randomly cut ∼25 1-2 cm long root segments. The full experimental setup had two geographic pairs (run separately for each pair) × two soil nutrient treatments × five replicates × 13 planting combinations comprising the various soil inoculum and competition treatments. However, we were only able to grow enough *P. tremuloides* seedlings of similar sizes for four replicates. This resulted in 234 total pots containing 360 total plants from which we made 288 plant measurements based on the plant competition × soil inoculum design (the two plant measurements in intraspecific competition treatments were averaged).

### Analysis

We analyzed *Populus* EM root colonization levels from the field and experiment to test Hypothesis 1. For field-collected roots, we created a linear model with *Populus* species, decomposition rate (projected as a measure of climatic influence on nutrient availability), and their interaction as fixed effects, and EM colonization as the response. We used multiple regression to assess the relative importance of predicted decomposition rates for explaining EM colonization compared to nine other soil and climatic variables (‘relaimpo’ package in R; Grömping 2006; R Core Team 2021; Supplemental Figure 1). For *Populus* root samples collected from the growth room experiment, we created a mixed effects model using the ‘nlme’ package (Pinheiro et al. 2021) with EM colonization as the response variable, inoculum type (i.e., *Populus, Acer*, or Sterilized inoculum), plant species (*P. deltoides* or *P. tremuloides*), N fertilization, competition type, and all interactions as fixed effects, and block as a random effect.

We compared plant growth responses between soil inoculum treatments to test Hypothesis 2. Our experiment used a Home vs. Away soil inoculum design: “Home” treatments refer to soil inoculum sources matched with the same plant species and “Away” treatments refer to the mismatch between plant species and soil inoculum source. Using only the single plant treatments (Figure 1B bottom row) we created a mixed effects model with aboveground biomass as the response (log transformed for normality), soil inoculum source (Home or Away), plant species, N fertilization, and their interactions as fixed effects, and block as a random effect. We calculated individual plant-soil feedback effects for each species as the log ratio growth response between Home vs. Away soil inoculum in each replicate. To characterize the net effect of soil communities on plant community dynamics, we calculated pairwise plant-soil feedback coefficients (*I*_*S*_) for each replicate as

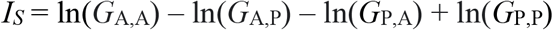

where *G*_i, j_ represents the biomass of plant i when grown in soil cultivated by plant j (i and j being A=*Acer* or P=*Populus*). Therefore, *G*_A,A_ and *G*_P,P_ are growth responses of *Acer* and *Populus* to their conspecific soil community, respectively, and *G*_A,P_ and *G*_P,A_ are growth responses of *Acer* and *Populus* to the heterospecific soil community, respectively (Bever 2003).

We tested Hypothesis 3 by examining how plant-soil interactions relate to species niche and fitness differences. Following Ke & Wan (2020), we used the full set of plant-soil and plant-plant treatments to calculate and partition interaction coefficients into different terms for plant competition or soil microbial effects. We calculated interaction coefficients using plant biomass measurements (averaged across soil replicates) when species *i* was grown with or without competition from species *j* in either Live or Sterilized soil inoculum treatments (denoted as *B*_*i,j,k*_, with *j* = 0 indicating no competitors and *k* = sterilized if soil inoculum was sterilized). These interaction coefficients reflect the per-capita effects of *j* on the biomass of *i* relative to the biomass of *i* when grown individually in sterilized soil. Microbial effects can be quantified by comparing interaction coefficients calculated from competition treatments using either Live or Sterilized soils of species *j*. Plant-plant interaction, in the presence of soil microbial effects, can be calculated as:

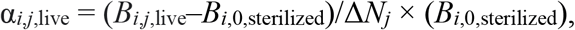

and in the absence of microbial effects as:

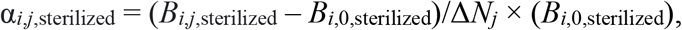

where *B*_*i,j*,live_ and *B*_*i,j*,sterilized_ represent the biomass of species *i* under the competitive impact of species *j* in Live and Sterilized inoculum of soil *j*, respectively, *B*_*i*,0,sterilized_ represents the biomass of species *i* grown individually in Sterilized inoculum, and Δ*N*_*j*_ represents the density of species *j* competitors (in this case, Δ*N*_*j*_ = 1). Here, it is important that the Live soil treatments are cultivated by the competitor to correctly account for their impact via both plant-plant competition and plant-soil interactions (Ke & Wan 2020). This approach was also used to calculate the relevant interaction coefficients from intraspecific competition pairs (i.e., *B*_*i,i,k*_ and *B*_*j,j,k*_). For each species pair and soil treatment combination, we quantified niche overlap (ρ, or the magnitude of niche difference, 1 – ρ) as the relative difference between interspecific and intraspecific interaction coefficients:

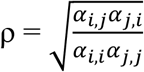

We quantified fitness ratios as:

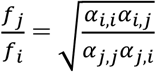

Under modern coexistence theory, competitive outcomes depend on two mechanisms related to niche differences and fitness ratios: stabilizing forces increase niche differences, while equalizing forces reduce fitness ratios.

## Results

Under natural conditions, *P. tremuloides* in colder, drier climates (higher latitude/elevation) formed 82% more EM root symbioses than *P. deltoides* in warmer, wetter habitats (lower latitude/elevation) (F_1,33_=14.9, p<0.001; Figure 2A). Projected litter decomposition rates explained 31% of the overall variation in EM colonization (F_1,33_=5.0, p=0.03) with a lower prevalence of EM symbioses in climates favoring faster nutrient turnover. These climate-constrained decomposition rates were the best predictor of EM colonization compared to other environmental variables (Supplemental Figure 1).

**Figure 2.**
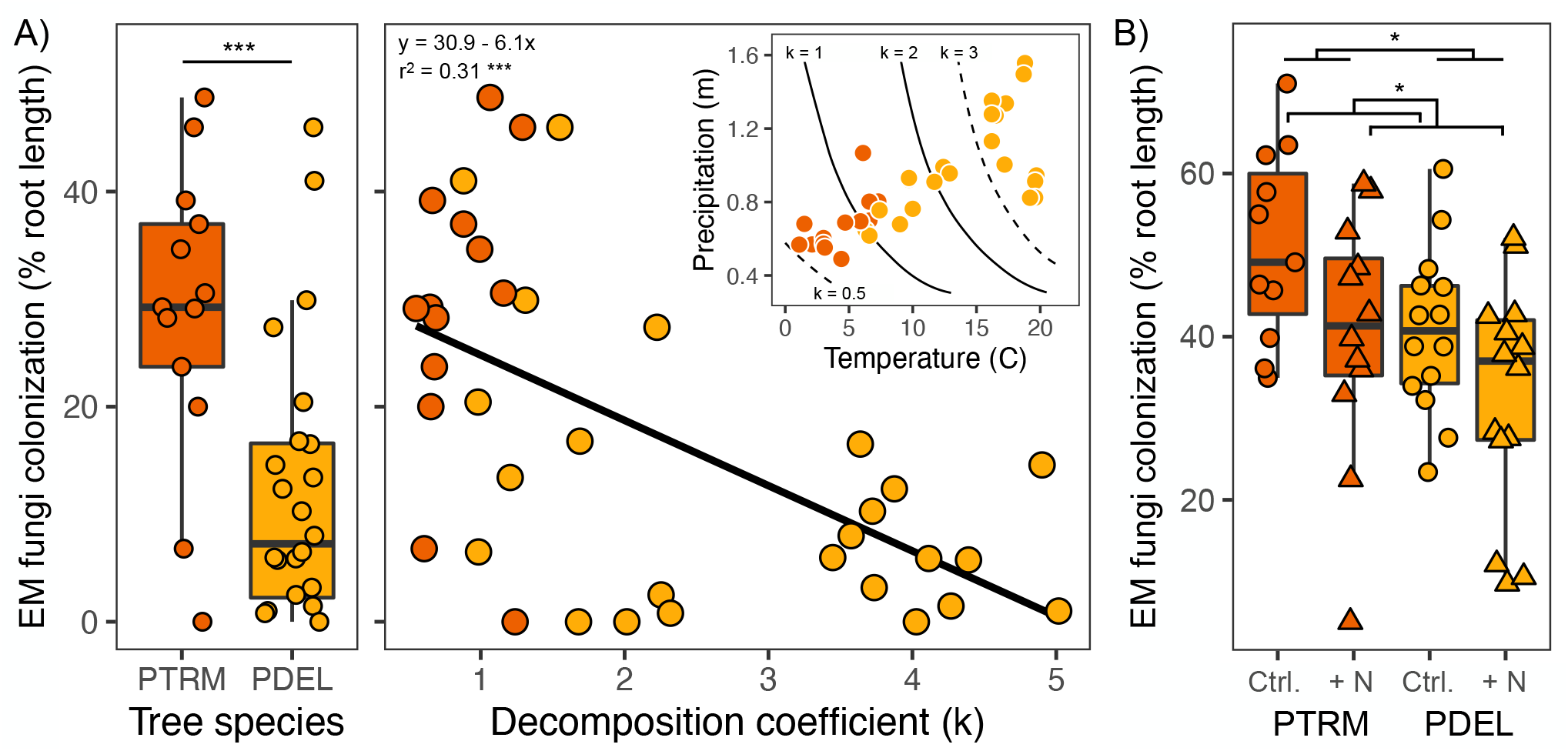
Pattens of ectomycorrhizal (EM) prevalence on *Populus* roots. **(A)** Field samples show that *P. deltoides* and *P. tremuloides* differ in the percent root length colonized by EM fungi overall, and EM colonization declines with higher decomposition coefficient values (k) that correspond with faster rates of nutrient turnover. The inset figure shows how field sites are located across temperature and precipitation gradients that control decomposition rates (lower k values reflect slower decomposition predicted by cooler, drier climates in the Yasso07 model; Tuomi et al. 2009). **(B)** In the experiment, EM root colonization varies between the two *Populus* in the same direction as field patterns (*P. tremuloides* > *P. deltoides*) and is reduced by inorganic nitrogen (N) fertilization in both species. Triangles show N fertilizer treatments, circles show unfertilized control treatments, PDEL = *P. deltoides*, PTRM = *P. tremuloides*.

In the growth room experiment, *P. tremuloides* showed 21% higher EM root colonization compared to *P. deltoides* with Live Home soil inoculum treatments (F_1,39_=6.6, p=0.01; Figure 2B), and there was no *Populus* species x fertilizer interaction (F_2,39_=0.3, p=0.6; Supplemental Table 1). Both *Populus* species had similar amounts of EM colonization when grown alone (∼36-41% root length) but showed different reactions to the competition treatments (species x competition interaction: F_2,39_=4.2, p=0.02). In their Home soil, EM colonization on *P. tremuloides* roots increased by more than a third in competition pairs – regardless of the competitor’s identity – compared to plants grown alone, while *P. deltoides* showed no changes in EM affinity when alone or in competition treatments (Supplemental Figure 2). As expected, the sterilized and *Acer* soil inoculum treatments resulted in very little to no EM root colonization for either *Populus* species (Supplemental Figure 3).

Experimental soils in the N addition treatment had 48% greater inorganic N supply rates compared to control soils (N addition mean=1488±151 μg N/10 cm^2^/8 weeks, Control mean=912±58 μg N/10 cm^2^/8 weeks). Adding mineral N also shifted the balance of other limiting nutrients (Supplemental Table 2), including soil phosphorus which was 112% lower in N addition treatments compared to control soils (N addition mean=2.9±0.2 μg P/10 cm^2^/8 weeks, Control mean=10.3±0.9 μg P/10 cm^2^/8 weeks). Fertilizer treatments decreased *Populus* EM colonization overall by 23% compared to control treatments (F_1,39_=7.4, p=0.01; Figure 2B).

Plant aboveground biomass varied by a significant interaction between plant species x soil inoculum source (Live Home vs. Away inoculum) (F_3,52_=8.0, p<0.001; Figure 3A, Supplemental Table 3). For the northern species, *P. tremuloides* grew better (Tukey’s post-hoc p<0.001) and *A. saccharum* performed worse (Tukey’s post-hoc p=0.01) in Home vs. Away soil treatments. This created a positive individual plant-soil feedback effect for *P. tremuloides* (mean=0.39±0.15 SE; t-test difference from zero: t_1,7_=2.6, p=0.02) and a marginally negative feedback effect for *A. saccharum* (mean=-0.26±0.16 SE; t_1,7_=1.6, p=0.07) (Figure 3B). Both southern species (*P. deltoides* and *A. saccharinum*) showed no growth differences between the reciprocal inoculum treatments (neutral individual feedbacks). However, calculating pairwise plant-soil feedback effects showed a negative *I*_*S*_ for the southern species pair that persisted in both N treatments (mean=-0.47±0.09 SE; t_1,9_=5.3, p<0.001; Figure 3C). In contrast, the northern species feedback *I*_*S*_ was slightly positive overall (mean=0.29±0.22 SE; t_1,7_=1.4, p=0.11) and changed with N addition: control treatments had positive *I*_*S*_ (mean=0.67±0.23; t_1,3_=3.0, p=0.03) but N fertilization caused neutral *I*_*S*_ (t_1,3_=0.4, p=0.4). Both *P. tremuloides* and *A. saccharum* responded negatively to the N fertilization treatments while *A. saccharinum* performed better with added N (species × N fertilization interaction: F_3,52_=8.1, p<0.001; Supplemental Table 3).

**Figure 3.**
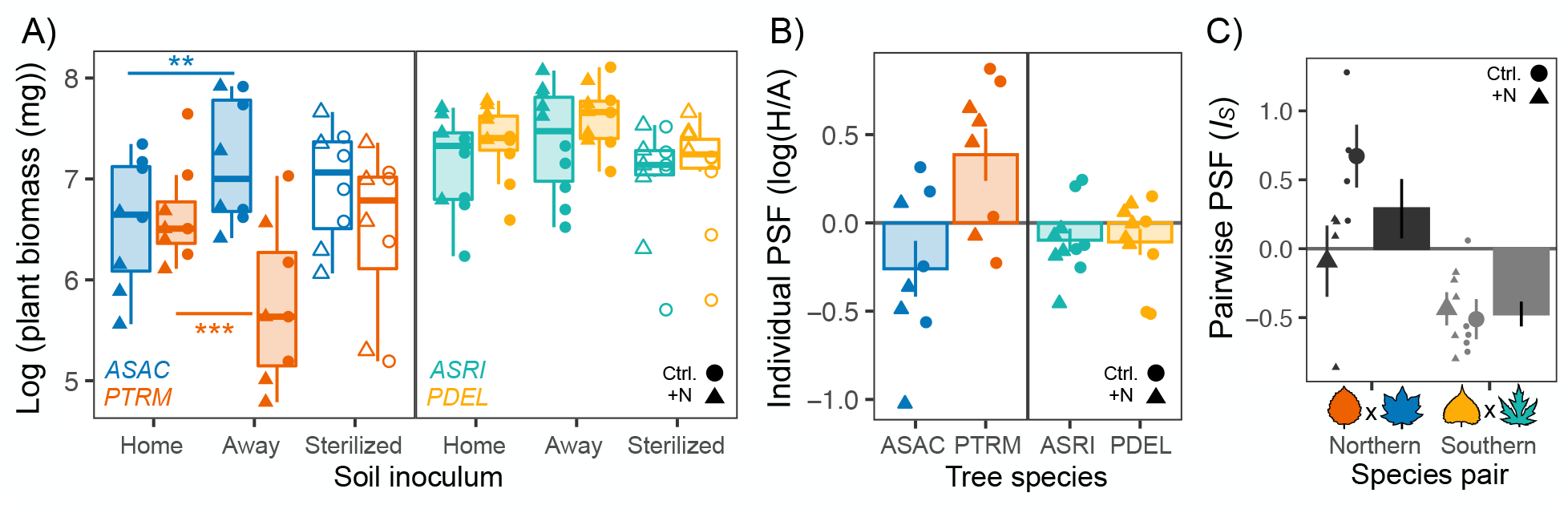
Soil microbial effects on plant growth and plant-soil feedbacks. **(A)** Aboveground biomass differences between Home vs. Away soil inoculum treatments show plant growth benefits of interacting with (*P. tremuloides*) or without (*A. saccharum*) their conditioned soil microbial communities. There we no clear growth differences between soil inocula for the Southern species pair. **(B)** Individual plant-soil feedback calculations for each replicate indicate stronger and more divergent effects in the Northern species pair (ASAC and PTRM) compared to the neutral effects in the Southern species pair. Bars show overall mean ± 1 SE. (**C**) Pairwise plant-soil feedbacks calculated for each species pair show differential responses to soil microbial communities result in negative coefficients that stabilize southern tree species coexistence (*I*_s_ < 0), while northern species interactions are moderately destabilized from positive plant-soil feedbacks (*I*_s_ > 0). Large points show nitrogen fertilization treatment means ± 1 SE, bars show overall mean ± 1 SE. Triangles show nitrogen fertilizer treatments, circles show unfertilized control treatments, ASAC = *A. saccharum*, ASRI = *A. saccharinum*, PDEL = *P. deltoides*, PTRM = *P. tremuloides*.

For the northern pair (*P. tremuloides* and *A. saccharum*), the net effect of soil microbes destabilized plant species interactions by causing niche differences to become increasingly negative (Table 1; Figure 4A). In the unfertilized control treatments, Sterilized inoculum predicts *A. saccharum* will win because of its stronger interspecific effect on *P. tremuloides* (α_P,A_). Incorporating Live soil microbes weakened the competitive impact imposed by *P. tremuloides* (α_P,P_ and α_A,P_). This caused interspecific competition (α_A,P_) to become slightly stronger than intraspecific competition (α_P,P_) for *P. tremuloides* in the more N-limited control soils, driving the interaction to be governed by priority effects where the first plant to arrive wins as neither species can successfully invade the other at their equilibrium (Figure 4A). With inorganic N fertilization, Sterilized treatments predicted stable coexistence, but adding Live soil microbes resulted in competitive dominance by *P. tremuloides*. Specifically, N fertilization in Sterilized treatments caused the strong interspecific competitive impacts of *A. saccharum* to become weaker (α_P,A_) and this led to coexistence (Figure 4A). However, adding mineral N in Live inoculum treatments caused *P. tremuloides* to inflict a stronger interspecific effect on *A. saccharum* (α_A,P_) and drove their competitive exclusion. We estimated the individual contribution of each plant species’ soil conditioning effects on the net competitive outcome by sequentially replacing each species interaction coefficients from Live treatments with the corresponding Sterilized coefficients and recalculating niche and fitness differences. Here, *P. tremuloides* soil microbes had the greatest influence on the competitive balance between the two northern tree species. Soil conditioning by *A. saccharum* either promoted their competitive dominance (low N) or maintained coexistence (high N), but strong destabilizing effects from *P. tremuloides* soil conditioning ultimately pushed the interaction towards priority effects (low N) or *P. tremuloides* competitive dominance (high N).

**Table 1.**
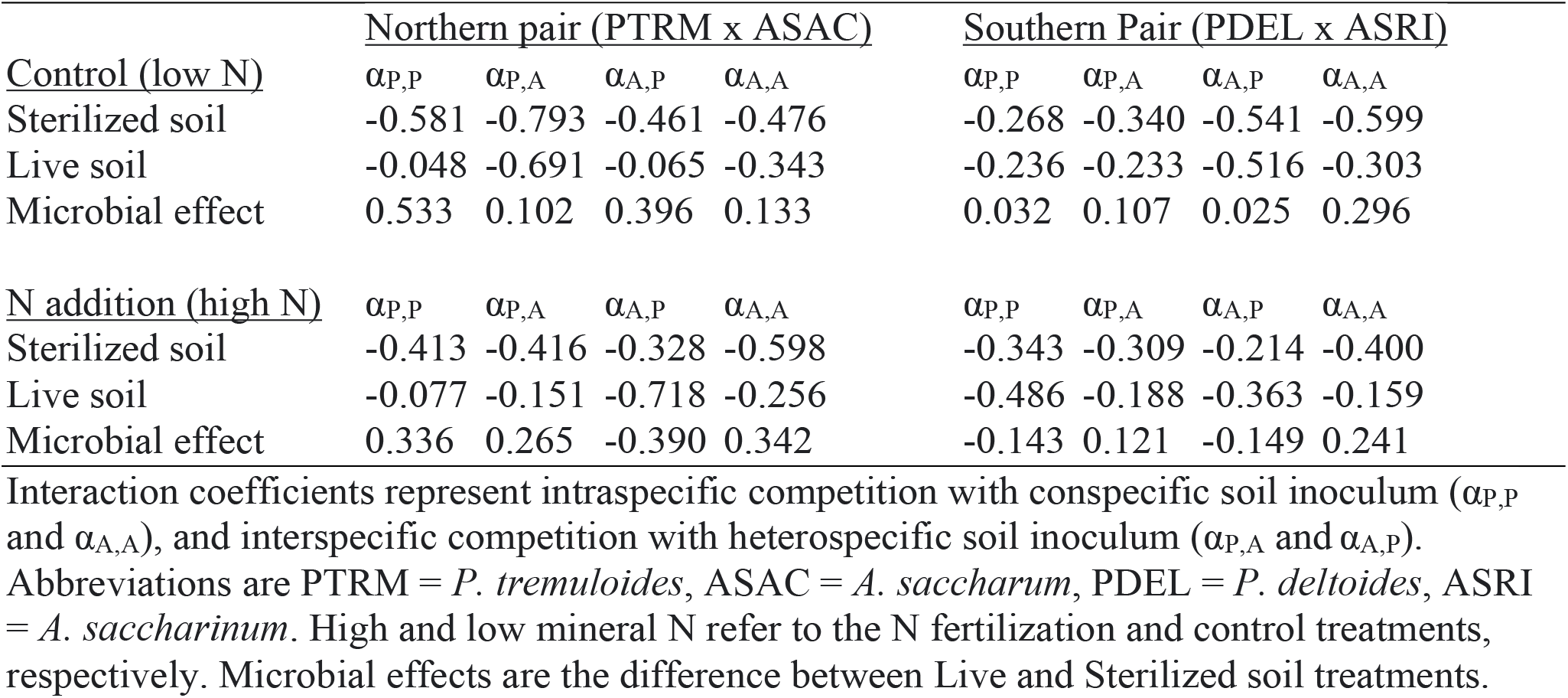
Interaction coefficients (α_ij_) of *Populus* and *Acer* tree species competition pairs.

|  | <u>Northern pair (PTRM x ASAC)</u> |  |  |  | <u>Southern Pair (PDEL x ASRI)</u> |  |  |  |
| --- | --- | --- | --- | --- | --- | --- | --- | --- |
| <u>Control (low N)</u> | $\alpha_{P,P}$ | $\alpha_{P,A}$ | $\alpha_{A,P}$ | $\alpha_{A,A}$ | $\alpha_{P,P}$ | $\alpha_{P,A}$ | $\alpha_{A,P}$ | $\alpha_{A,A}$ |
| Sterilized soil | -0.581 | -0.793 | -0.461 | -0.476 | -0.268 | -0.340 | -0.541 | -0.599 |
| Live soil | -0.048 | -0.691 | -0.065 | -0.343 | -0.236 | -0.233 | -0.516 | -0.303 |
| Microbial effect | 0.533 | 0.102 | 0.396 | 0.133 | 0.032 | 0.107 | 0.025 | 0.296 |
| <u>N addition (high N)</u> | $\alpha_{P,P}$ | $\alpha_{P,A}$ | $\alpha_{A,P}$ | $\alpha_{A,A}$ | $\alpha_{P,P}$ | $\alpha_{P,A}$ | $\alpha_{A,P}$ | $\alpha_{A,A}$ |
| Sterilized soil | -0.413 | -0.416 | -0.328 | -0.598 | -0.343 | -0.309 | -0.214 | -0.400 |
| Live soil | -0.077 | -0.151 | -0.718 | -0.256 | -0.486 | -0.188 | -0.363 | -0.159 |
| Microbial effect | 0.336 | 0.265 | -0.390 | 0.342 | -0.143 | 0.121 | -0.149 | 0.241 |

Interaction coefficients represent intraspecific competition with conspecific soil inoculum ( $\alpha_{P,P}$ and $\alpha_{A,A}$ ), and interspecific competition with heterospecific soil inoculum ( $\alpha_{P,A}$ and $\alpha_{A,P}$ ). Abbreviations are PTRM = *P. tremuloides*, ASAC = *A. saccharum*, PDEL = *P. deltoides*, ASRI = *A. saccharinum*. High and low mineral N refer to the N fertilization and control treatments, respectively. Microbial effects are the difference between Live and Sterilized soil treatments.

**Figure 4.**
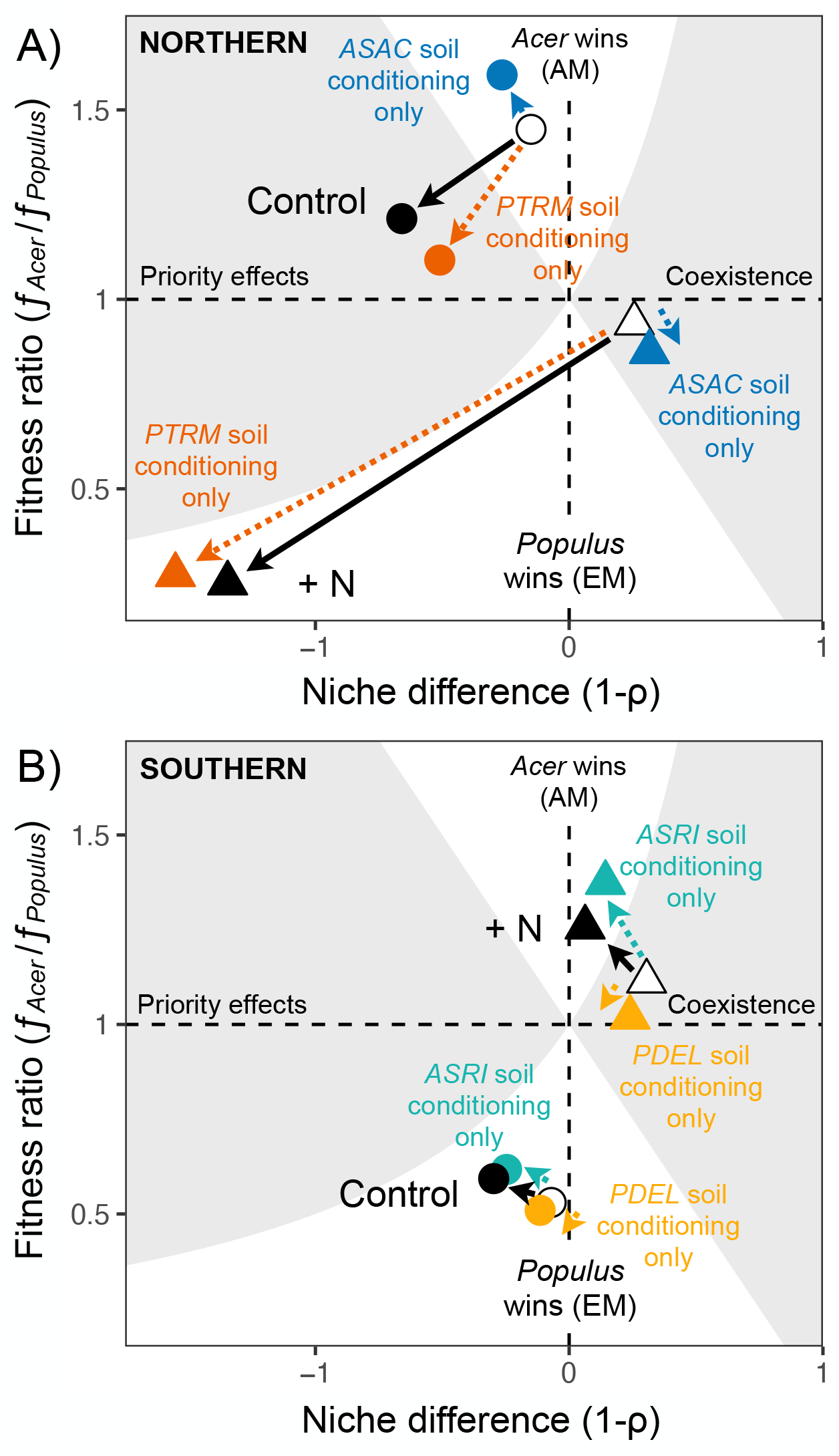
Soil microbial effects on the outcomes of plant competition under different soil nutrient conditions. The parameter space shows the niche difference (i.e., stabilization potential, x-axis) and fitness ratio (y-axis) of the Northern **(A)** and Southern **(B)** plant species interactions. Shaded areas indicate regions where coexistence (right) or priority effects (left) are expected to occur, and unshaded regions show competitive exclusion by *Acer* (top) or *Populus* (bottom). Circles show Control (unfertilized) treatments and triangles show inorganic N fertilization treatments. Open and closed symbols indicate Sterilized and Live soil inoculum, respectively, with solid arrows showing the net effect of soil microbes on plant competitive outcomes. Dashed arrows show the influence of species-specific soil conditioning effects on plant competition (different colors). Circles show unfertilized control treatments, and triangles show nitrogen fertilizer treatments.

The net effect of soil microbes on the southern tree species competition pair (*P. deltoides* and *A. saccharinum*) was weaker compared to northern species overall (Table 1; Figure 4B). In low N control treatments, *P. deltoides* showed competitive dominance over the co-occurring *A. saccharinum* regardless of soil microbial effects from either species. Specifically, *P. deltoides* wins under low N in Sterilized treatments because of its stronger interspecific impact on *A. saccharinum* (α_A,P_), and the relative strength of intraspecific and interspecific competition showed little change with Live microbes such that the competitive outcome remained the same. As with the northern pair, N fertilization enabled coexistence between the southern species pair in Sterilized inoculum treatments. This was caused by N addition weakening the otherwise strong interspecific competitive impact of *P. deltoides* on *A. saccharinum* (α_A,P_). With Live inoculum, N fertilization reduced *A. saccharinum* intraspecific effects (α_A,A_) resulting in the competitive exclusion of *P. deltoides* as *A. saccharinum* microbial conditioning effects exacerbate species fitness differences under higher N conditions.

## Discussion

Studies of geographic variation in root symbioses tend to treat plant mycorrhizal type as monolithic functional classes (Soudzilovskaia et al. 2015; Bueno et al. 2017; Lu & Hedin 2019; Stedinger et al. 2019). However, our results show that the extent of mycorrhizal associations within a functional class can also show large geographic changes, and that these within group patterns match expectations from studies comparing mycorrhizal nutrient uptake strategies. Previous work has shown that *Populus* can vary in the degree of EM and AM symbioses (Gehring et al. 2006; Karst et al. 2021a), and here we demonstrate why such patterns might arise based on geographic differences in mycorrhizal strategies and soil nutrient availability. In support of Hypothesis 1, the more northern species (*P. tremuloides*) had greater EM colonization than the more southern species (*P. deltoides*), aligning with global patterns of increasing EM dominance with latitude and the climate controls on nutrient cycling (Steidinger et al. 2019). Although we did not measure AM root colonization in this study, *P. tremuloides* has generally low AM affinity (Karst et al. 2021a) while *P. deltoides* shows appreciable AM colonization (Khasa et al. 2002). Stronger soil N-limitation favors EM symbioses to become more prevalent as plants increasingly rely on these fungi to mine nutrients from organic sources (Lu & Hedin 2019), and here we found evidence supporting this within a single plant genus across geographic scales. Our experimental results support this further as N addition reduced *Populus* EM colonization levels, consistent with past work demonstrating the negative effects of nutrient fertilization on mycorrhizal associations, symbiotic benefits, and EM community composition (Johnson 1993; Morrison et al. 2016; van der Linde et al. 2018; but see Karst et al. 2021b).

There is likely a limit on how flexible plants can be regarding the intensity of mycorrhizal colonization. For instance, *P. deltoides* lacks specific genetic loci found in other poplars that promote the recognition and formation of EM symbioses with *Laccaria bicolor* (Labbé et al. 2019). If this extends to other fungi, it could be another reason why *P. deltoides* had lower EM affinity. Such inflexibility might occur in the opposite direction too if highly EM plants are unable to advantageously lower their symbiotic associations down to a more cost-effective level under weak nutrient-limitation. The decline of EM trees with N pollution across US forests might be one potential example of this phenomena (Averill et al. 2018). Even if plants can reduce EM colonization slightly (as found here), it may not be enough to remain competitively viable in communities with differing plant-mycorrhizal strategies. This raises questions about the functional consequences of variation in mycorrhizal dependence across plant species or genotypes as soil resources change.

Diverse mycorrhizal strategies have been shown to influence the mechanisms underlying plant coexistence by affecting niche partitioning and relative fitness differences (van der Heijden et al. 2003, 2008; Stanescu & Maherali 2017; Peay 2018), but the nature and design of these studies made them largely incapable of predicting whether plant-soil interactions lead to stable coexistence or species exclusion. Our results demonstrate the usefulness of modern coexistence theory to understand how soil microbiome and nutrient variation interactively shape the outcomes of plant-plant competition. We find that competitive outcomes between plants with different mycorrhizal fungi is contingent on their geographic origin and the soil nutrient context. Adding soil microbes did not promote stable coexistence among AM and EM tree species – Live inoculum always had destabilizing effects and sometimes contributed to competitive exclusion by intensifying fitness differences. This strong tendency for microbially-driven competitive exclusion among tree species may help explain the bimodal signature in models of EM vs. AM competitive strategies (Lu & Hedin 2019) and the abrupt latitudinal transitions in forest mycorrhizal states (Steidinger et al. 2019). Although we did not directly manipulate mycorrhizal fungi in the experiment, our whole microbiome inoculum approach mirrors how plants compete in nature by recruiting into competitor soils, which clearly affected EM colonization rates.

Most conceptualizations of the functional differences between AM and EM symbioses focus on nutrient competition (Phillips et al. 2013; Corrales et al. 2016; Averill et al. 2019), and we find evidence that inorganic N fertilization switches AM vs. EM competitive outcomes (often in the expected direction). In Sterilized inoculum, *Populus* and *Acer* have enough niche separation to coexist with high N availability. However, incorporating soil microbiome effects for the southern species pair matches the AM-EM nutrient competition framework (support for Hypothesis 3): *A. saccharinum* and *P. deltoides* were superior competitors in high and low mineral N conditions, respectively. Higher latitude soils are typically characterized as N-limited (Du et al. 2020), meaning that the competitive outcomes between northern species in unfertilized control treatments are more likely to reflect the actual soil nutrient properties in the field. Under these conditions, *P. tremuloides*-cultivated microbes pushed the interaction to be governed by priority effects, which allow for alternative stable states depending on the initial species density. This exactly fits the bimodal patterns that emerge from evolutionary stable state models of plant-mycorrhizal nutrient economies combined with largescale observations of AM and EM tree species prevalence across US forests (Lu & Hedin 2019). Overall, soil microbiome effects weakened the competitive impact imposed by *P. tremuloides*, and we speculate that EM symbioses formed in Live inoculum gave *P. tremuloides* greater access to the organic nutrient pool, thereby relaxing resource competition with competitors.

However, *P. tremuloides* does not completely fit the plant-mycorrhizal nutrient competition paradigm as this northern, highly EM species was projected to be competitively superior with greater inorganic N levels. Some evidence indicates EM fungi may have the potential to hoard N (Nasholm et al. 2013; Franklin et al. 2014), meaning that *P. tremuloides* could have captured a competitive advantage by monopolizing soil resources in the fertilization treatments. This unexpected pattern may also be related to the controlled conditions in which the interactions took place (i.e., missing an important detail from natural systems or the methods for establishing plants in the experiment), or that P limitation played some role worth exploring in the future. We likely do not fully understand how mycorrhizal fungi affect nutrient competition, and this unexpected outcome reinforces the need for mechanistic experiments between mycorrhizal associations.

We used multiple approaches to characterize soil microbial effects on plant coexistence, from which there were both complementary and contradictory lines of evidence. The plant-soil feedback and modern coexistence frameworks converged on the same pattern for the northern species pair. Specifically, positive plant-soil feedbacks (pairwise *I*_*S*_ > 0) should lead to positive frequency dependence and priority effects (dominance of either species). This was confirmed in the modern coexistence theory approach as adding *P. tremuloides* soil microbes destabilized the overall interaction leading to priority effects (low N) and the competitive exclusion of *A. saccharum* (high N). Plant-soil interactions in the southern species pair were predicted to stabilize coexistence with net negative pairwise feedback effects (*I*_*S*_ < 0). However, analyzing their interaction coefficients in a modern coexistence framework showed destabilizing effects with the two species never predicted to coexist and, instead, that they should competitively exclude one another under different soil nutrient conditions.

The mismatch between plant-soil feedback direction and the projected competitive outcome arises because the *I*_*S*_ pairwise feedback index uses plant biomass measurements when individuals were grown alone in different soils, whereas the other interaction coefficient calculations incorporate both plant-soil and plant-plant effects. Even though both southern species performed slightly worse when grown alone in their Home inoculum, each species benefited from conspecific soil conditioning in a net competitive sense across the N treatments (e.g., mostly positive *Microbial effects* in Table 1). In other words, microbial effects ameliorated competition for the resident plant species more than for the invader. Thus, the integrated approach captures an element of positive microbial effects that is missed by traditional plant-soil feedback calculations. In fact, only the modern coexistence approach gave predictions that matched our intuition – a shift towards AM competitive dominance with higher N in the southern pair. Such discrepancies underscore that the plant-soil feedback approach alone may be insufficient to accurately predict coexistence outcomes without explicitly incorporating plant-plant competition (Ke & Wan 2020). Collectively, our findings empirically support emerging theoretical work on how mutualisms alter species coexistence and fit largescale observations of mycorrhizal biogeography across resource gradients, further emphasizing the importance of incorporating plant-mycorrhizal associations into the mechanisms of biodiversity maintenance.

## Acknowledgments

This research was funded in part by the US Department of Energy Biological and Environmental Research program Award DESC0016097 to KGP and the Yushan scholar program of Taiwan MOE (NTU-110VV010) to PJK. We thank Caroline Daws for field assistance and Claire Willing and Cong Wang for growth room support. The authors declare no conflicts of interest.

